## Supplementary Figures and legends for "An upstream secondary DNA motif within the *IL3* insulator CTCF binding site is required for enhancer-blocking insulator activity"

S1 figure **CTCF sites at me3K27H3 chromatin boundaries**

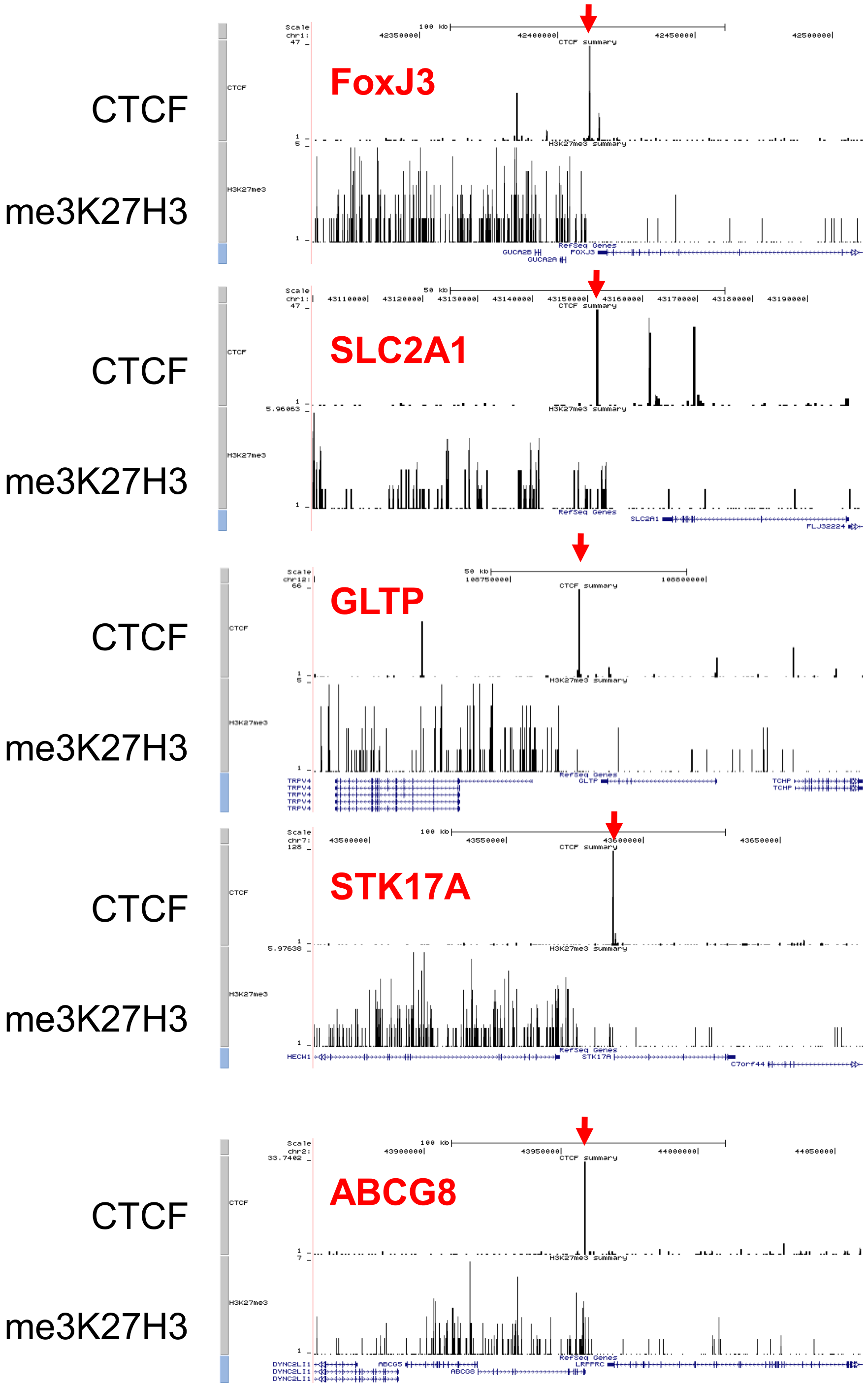


**S1 Figure. Locations of CTCF sites at histone H3K27me27 chromatin boundaries.**
Data was extracted from a UCSC genome browser session for which the authors had uploaded their published histone H3K27me27 and CTCF ChIP data. This session was accessed via a link in the authors publication: https://dir.nhlbi.nih.gov/papers/lmi/epigenomes/hgtcell.aspx [24, 47].

### S2 figure Unmanipulated original images used in figure 2

The boxed regions of panel C indicate irrelevant lanes that were removed before preparing Fig 1C.

Fig 2C original image with deleted lanes marked by boxes

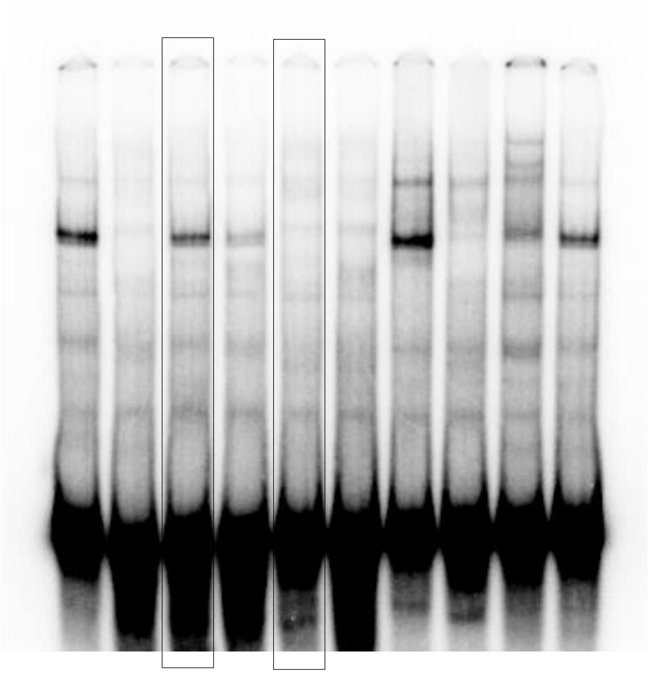

Fig 2E original image

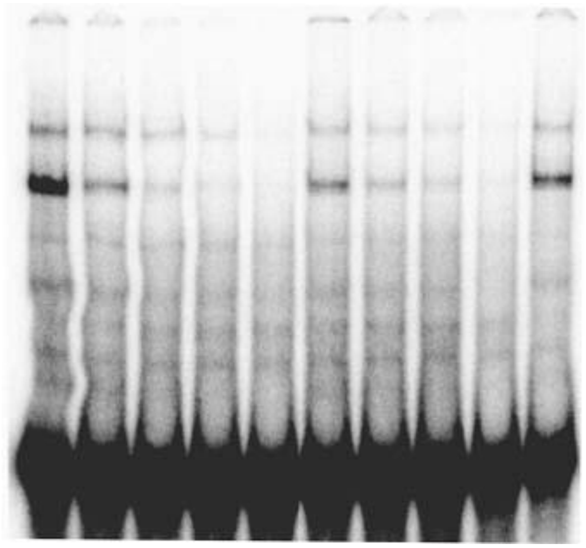

Fig 2F original image

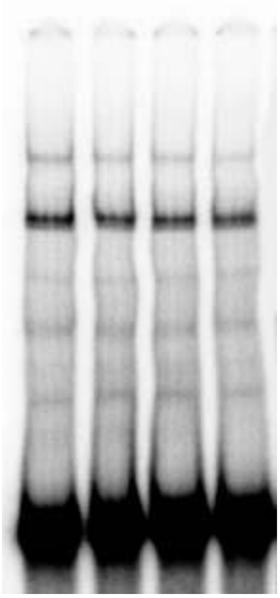

### S3 Table CTCF sites in the IL-3 gene insulator

```
                       2° upstream          1° core
                          motif              motif
Global consensus       CTGCAGTTCCNNNNNTGGCCACCAGGGGGCGCC
of composite sites     G  TTACGTT     C     G   AT   AGT
                            C

IL-3 +2.9 kb
HUMAN       AGGGGCGGCGGCTGCAGTTTCTGGAAGGACCACTAGGGGGAGACATGC
ORANGUTAN   ......................................A.........
RHESUS      .........T..............................G...G...
BABOON      .........T..................................G...
MOUSE LEMUR .....AC..T.....G.Gg..C..C......t.GG....AG.cGTGAG
BUSHBABY    ....AGT..T................t....t....a......G....
TREE SHREW  T....AT..A.........c......A............A..g..G..
MOUSE       TA...ACA.T.........ctAA.T.t...............g...T.
RAT         TA.A.GCA.T.........ctAA.T.t..........A..GAg...T.
GUINEA PIG  T..A.AT..TA............A..t.............cAgtG.C.
RABBIT      GAAA..T..T.....t  ..........A.........a.....G...T.
HORSE       T.A..AT..T...........CA.TGtAG.............g.....
DOG         .A.A.AA..TC.............T..A.............Ag..G..
ELEPHANT    TA..AACCTT.g..t.........T...G...................
ROCK HYRAX  ---------A.g............T...G..t...A..........A.
ARMADILLO   GA...AA..T.............TC.A.G....C.C...A..g.....
SLOTH       TA...AT..T.............TCT..G....C........g.....


IL-3 +4.2 kb
HUMAN       AGAGGATGCTGCTGCAGTTTCTGGATTGACCACTAGGGGGAGGTGTGT
ORANGUTAN   ....................................a...........
RHESUS      .......................A........g...............
MARMOSET    ......G..............C..............a...........
MOUSE LEMUR TAG...CA...........c....TGG......c......c....CCG
BUSHBABY    TAG..CCTG..........c..TCTAA......c.............C
TREESHREW   T.G..C..TG.A.........CA..AG......a...A..c......C
MOUSE       CAGA.T.CT.T........cG..A.A..........a..A..AAAATG
RAT         .TGA..CAT.C........ct..A.A..........a..Ac.A.A..C
GUINEA PIG  CT........Ag...t......AACA.......c..a.A...c....C
ALPACA      CAG................c..T..GGAG..........AG.cACGCC
HORSE       T....................CA.TG.AG..............CA..C
CAT         T.GA..A.................TGGAG..........C...CAG.C
DOG         .AGA..A...C.............TAGA.............a.CCACC

Human IL-3 +2.9/+4.2 identities
            AG*GG**GC*GCTGCAGTTTCTGGA**GACCACTAGGGGGAG***TG*
```

**S3 Table. Conservation of the *IL3* insulator CTCF sites in mammalian species.**
Alignments of the conserved bases within the *IL3* +2.9 and +4.2 CTCF binding sites. The composite consensus sequence is shown above the sequence alignments. Identities are depicted as dots. Lower case letters within the alignment indicate an allowable base based on the consensus. Capital letters represent unfavourable sequence changes. Underlined segments represent conserved sequences.

### S4 Table UPK2 gene intron CTCF site

| | |
|---|---|
| | **2° upstream motif** **1° core motif** |
| **Global consensus of composite sites** | **CTGCAGTTCC**NNN**TGGCCACCAGGGGGCGCC** |
|  | G TTACGTT C G AT AGT |
| | C |
| HUMAN | GGTGCAGCTT**CTGCAGCCCA**AGCCTG**CCACCTGGTGGTCAT**ACT |
| CHIMP | ...........................................C |
| GORILLA | ...........................................C |
| ORANGUTAN | ................................A........... |
| RHESUS | ...........................................C |
| BABOON | ...........................................C |
| MARMOSET | CA...........................T..........c.TG |
| TARSIER | T.......................................c.GC |
| MOUSE LEMUR | T.......................................c..G |
| BUSHBABY | T.....CT....-...........................c... |
| MOUSE | TACC..C.A........t...........T..........G.TC |
| RAT | TACC..C.A........t...........T............TC |
| KANGAROO RAT | TACCT........t........T.................G... |
| GUINEA PIG | C.GT.......A....tg..........C........c..G... |
| SQUIRREL | TA............G.........................GG.. |
| RABBIT | CACC...T..........................C..G..G..C |
| PIKA | CA.C..............................C..G..G..A |
| DOLPHIN | CACCT............t.....................gc..A |
| COW | TACC.......................................G |
| HORSE | TA.C.G..................................G..A |
| CAT | .ACC.G....T...GT........................c..A |
| DOG | C..C....................................c..A |
| MICROBAT | TACC................C.............C....gG..C |
| MEGABAT | TA.C..................................TgG..C |
| HEDGEHOG | TA.T..............t.........C...........G..C |
| SHREW | .AC...........G...tG..G.....C........c..GG.C |
| ELEPHANT | TA.C.............................C...c..G..C |
| ROCK HYRAX | T.CC................................Ac..G..C |
| TENREC | TA.C....G.gG.....t.........................C |
| ARMADILLO | CCCC.G...............................c..G... |

**S4 Table. Conservation of the *UPK2* intronic CTCF site in mammalian species.**
Alignments of the conserved bases within the *UPK2* CTCF binding sites. The composite consensus sequence is shown above the sequence alignments. Identities are depicted as dots. Lower case letters within the alignment indicate an allowable base based on the consensus. Capital letters represent unfavourable sequence changes. Underlined segments represent conserved sequences.

### S5 Table HOXA9 chromatin boundary

```
                              2° upstream         1° core
                                 motif            motif
Global consensus               CTGCAGTTCCNNNTGGCCACCAGGGGGCGCC
of composite sites             G  TTACGTT   C     G   AT   AGT
                                    C

HUMAN                   GCTGAGGCTGCAGTACCAAACGGCGGCCAGCAGATGGCAGTG
CHIMP                   ...................G......................
GORILLA                 ..........................................
ORANGUTAN               .......A.........G..TA.T..................
RHESUS                  ......A..........G..T.AT..................
SQUIRREL                T.G.G.........G.T..GGTAA......g...........
MOUSE                   ....C.........G....GAAAATT....g...........
RAT                     ....C.........Gtt..GAAAA.T....g...........
GUINEA PIG              ..C.G.........G.GCGCGC..A..............C..
RABBIT                  --------......Gt.C.GGCC...................
PIKA                    --------......G..C.G.AT...................
DOLPHIN                 ...TG-........Gt..GGGT...................A
COW                     ....G-........Gt...GGCA..................A
CAT                     ....G.........G..C.GATT..................A
DOG                     ..C.G.........G.AG.GAT....................
HEDGEHOG                ..G.G-A.......G..GGCAA.T.A....a...........
SHREW                   ..G.--.TC.....G.GGCG......................
ELEPHANT                --CTG-........G..TC.GT....................
TENREC                  --CTGC...........GC.GTTT..T..........G..CA
ARMADILLO               --G.GA........T..T.CG.AT..................
```

**S5 Table. Conservation of the *HOXA9* chromatin boundary CTCF site in mammalian species.** Alignments of the conserved bases within the *HOXA9* CTCF binding sites. The composite consensus sequence is shown above the sequence alignments. Identities are depicted as dots. Lower case letters within the alignment indicate an allowable base based on the consensus. Capital letters represent unfavourable sequence changes. Underlined segments represent conserved sequences.

**S6 table CTCF sites at histone H3 K4me3 chromatin boundaries**

### FoxJ3 Gene

```
>hg18_dna range=chr1:42411751-42411798
CGGCAGCCCCGCCCCCTAGTGGCCACAGGCACTCACTACAGCCTAATT

                       2° upstream          1° core
                          motif              motif
Global consensus       CTGCAGTTCCNNNNNNTGGCCACCAGGGGGCGCC
of composite sites     G  TTACGTT      C     G   AT   AGT
                            C
HUMAN           AATTAGGCTGTAGTGAGTGCCTGTGGCCACTAGGGGGCGGGGCTG
RHESUS          ...G......c......................a...T.......
TREE SHREW      .G.GG.....c......CA...A.......c..a.......AG..
MOUSE           .GGC......c.TG..A........AT...c.......a.cC...
RAT             GCAA......c.TG..A.........T...c.......accC...
GUINEA PIG      ..AG......ct................g.c..a....a.AT...
RABBIT          CG.G......c...A..CA......A....c..a....a......
HEDGEHOG        GC.C......c..-...CA.TA.......gc..a....a.A.GGC
DOG             .GCG......c......C.G.A.c....ggc..a.....c.TGGC
CAT             .GCG......c......CA..G.G......c..a....a.A.GGC
HORSE           .G.G......c......CA..G.A......c..a......A.GGC
COW             C..G......c...A..GAG.A.A....g.c..a.....cA.GGC
ARMADILLO       ...GT.....c......CA.....t.....c..a....a.A.GGC
ELEPHANT        ...G.A....c..CA..CA......A....c..a......A.GGC
TENREC          CG.G.A....c......CA...........c..a..Ct.AcTGCT
```

### ABCG8 gene

```
>hg18_dna range=chr2:43958688-43958741
CAGGGTGCTGCAGTGGCACAGACCAGCCACAGGATGGCAGTAGAATAAAGACAGTCGAAAG

               2° upstream          1° core
                  motif              motif
Consensus      CTGCAGTTCCNNNNNNTGGCCACCAGGGGGCGCC
               G  TTACGTT      C     G   AT   AGT
                    C
HUMAN       GTGCTGCAGTGGCACAGACCAGCCACAGGATGGCAGTAGAATAAAGACAGTCGA
RHESUS      ......................................................
TREE SHREW  .C......c....C......T..............................T.G
MOUSE       .................GT................................T..
RAT         .................GT................................T..
GUINEA PIG  .................GT................................T..
RABBIT      .G..........t....GT.G..............................T..
DOG         A..T.........G...G.TG..............................A..
CAT         AATg.............G.TG..............................G..
HORSE       .G...A...........G..G................................G
COW         C................G.TG...............................A.
ELEPHANT    .G............G.AGT.G.....G........................A..
TENREC      CG......c.......C...G.............................AT..
```

**S6 Table. Conservation of chromatin boundary CTCF sites in mammalian species.**

Alignments of the conserved bases within the *FOXJ3* and *ABCG8* CTCF binding sites. The composite consensus sequence is shown above the sequence alignments. Identities are depicted as dots. Lower case letters within the alignment indicate an allowable base based on the consensus. Capital letters represent unfavourable sequence changes. Underlined segments represent conserved sequences.
